# A novel panel of short mononucleotide repeats linked to informative polymorphisms enabling effective high volume low cost discrimination between mismatch repair deficient and proficient tumours

**DOI:** 10.1101/305383

**Authors:** Lisa Redford, Ghanim Alhilal, Stephanie Needham, Ottie O’Brien, Julie Coaker, John Tyson, Leonardo Maldaner Amorim, Iona Middleton, Harsh Sheth, Osagi Izuogu, Mark Arends, Anca Oniscu, Ángel Miguel Alonso, Sira Moreno Laguna, Mauro Santibanez-Koref, Michael S. Jackson, John Burn

## Abstract

Somatic mutations in mononucleotide repeats are commonly used to assess the mismatch repair status of tumours. Current tests focus on repeats with a length above 15bp, which tend to be somatically more unstable than shorter ones. These longer repeats also have a substantially higher PCR error rate and tests that use capillary electrophoresis for fragment size analysis often require expert interpretation. In this communication, we present a panel of 17 short repeats (length 7–12bp) for sequence-based microsatellite instability (MSI) testing. Using a simple scoring procedure that incorporates the allelic distribution of the mutant repeats, and analysis of two cohort of tumours totalling 209 samples, we show that this panel is able to discriminate between MMR proficient and deficient tumours, even when constitutional DNA is not available. In the training cohort, the method achieved 100% concordance with fragment analysis, while in the testing cohort, 4 discordant samples were observed (corresponding to 97% concordance). Of these, 2 showed discrepancies between fragment analysis and immunohistochemistry and one was reclassified after re-testing using fragment analysis. These results indicate that our approach offers the option of a reliable, scalable routine test for MSI.

## Introduction

Two decades ago, Ionov *et al.* [1] reported that length variation in polyA mononucleotide repeats (MNR) was present in up to 12% of colorectal cancers (CRC), and was associated with a distinct pathological and molecular phenotype. Increased instability was observed for other microsatellites and this led to the designation of such tumours as microsatellite unstable. Microsatellite instability is a hallmark of tumours where the function of the mismatch repair (MMR) system is compromised [2]. MMR defects of clinical significance affect the *MSH2, MLH1, MSH6* and *PMS2* genes [3, 4]. Germline defects lead to an inherited autosomal dominant cancer predisposition syndrome, the Lynch Syndrome, that is characterised by a high risk for colon and endometrial cancer and an increased susceptibility to a variety of other malignancies [5]. The mismatch repair status of a cancer can be of clinical interest. Compared to other colorectal tumours, microsatellite unstable CRCs have been found to have a better prognosis [6]. More recently, patients with MMR deficient CRCs have been shown to benefit from immunotherapy with pembrolizumab [7]. Identification of patients with Lynch Syndrome also allows at-risk relatives to be identified [8]. This enables the implementation of monitoring regimes designed to detect cancer early, and the use of prophylactic measures such as the regular intake of aspirin, which has been shown to reduce cancer rates in patients with Lynch Syndrome by over 50% [9]. The implications for cancer treatment and management have led to recommendations to increase the proportion of CRC and endometrial tumours that are tested for MMR defects [10–13].

Testing for microsatellite instability (MSI) is one of the main methods used to assess MMR proficiency. However, somatic microsatellite mutations can also be observed in MMR proficient tumours. Thus, detection of low levels of microsatellites instability is not considered to be indicative of mismatch repair defects [14, 15]. MSI is commonly tested by amplification of a panel of microsatellites followed by analysis of the amplified fragments by capillary electrophoresis. A variety of panels have been recommended and current tests rely on long MNRs [16]. Long homopolymers tend to be more unstable both *in vivo* and *in vitro*, and PCR-induced errors lead to stutter peaks in electropherograms [17]. This can complicate downstream phenotype interpretation and visual inspection of the fragment size profiles can be required. Samples can be classified according to the frequency of microsatellite mutations. For example, the Revised Bethesda Guidelines for Hereditary Nonpolyposis Colorectal Cancer (Lynch Syndrome) and Microsatellite Instability described a classification using a panel of 5 quasi monomorphic MNR [18]. Samples showing mutations in two or more MMR designated as microsatellite instability high (MSI-H) samples, samples with only one altered MNR as microsatellite instability low (MSI-L) and where all microsatellites appear to be stable as microsatellite stable (MSS). MSI-H status is indicative of an MMR defect.

Microsatellite instability assesses the function of the MMR system. An alternative is to ascertain the presence of its components by immunohistochemistry (IHC). Lack of protein can result from mutations causing premature truncation of the encoded polypeptides and nonsense-mediated decay, or from the destabilisation of protein complexes leading to accelerated degradation of their components [19]. IHC requires highly skilled personnel. Since IHC assesses the levels of MMR proteins as opposed to a consequence of MMR dysfunction, there is some discordance between the results of microsatellite instability and IHC analyses [19, 20]. The reported concordance varies but a sensitivity of 92% for IHC in predicting MSI has been reported [19].

Several groups have developed sequencing based approaches to identify microsatellite instability. These include methods utilising genome [21] or transcriptome [22] wide data, as well as sequences from target enriched libraries [23, 24] and more recently, melt-curve analysis based test [25]. *In vitro* amplification errors, which lead to the presence of variant read lengths in the PCR product, can complicate sequence-based approaches. The frequency of such artefacts will differ between MNRs, but some mutant reads are expected even in the absence of mutations in the starting material. One approach to address the problem of amplification errors is to use a threshold value of the proportion of mutant molecules to discriminate between PCR-artefacts and the genuine presence of MNR mutations in the starting material [24].

Short MNRs tend to be less polymorphic than longer ones [26]. Thus, the likelihood of encountering germline variants in short MNRs is reduced, suggesting that they would be suitable for assessing MSI status in tumours without requiring matched germline DNA. The lower mutation rate also means that mutant reads from shorter repeats are more likely to reflect a single mutational event and affect only one allele while recurrent artefacts will affect both alleles. As a result, assessing whether length variants are concentrated in one allele offers an additional criterion to differentiate between PCR artefacts and mutations that occur *in vivo*.

The aim of this study was to develop a method suitable for high throughput and automated microsatellite analysis that allows separation of samples into two classes: those with clinically significant instability designated MSI-H and those with little or no evidence of instability and designated stable or MSS. The separation of unstable samples into “high” or “low” would thus be made redundant.

This involved selecting a panel of short MNR, and developing of a method to score instability based on both MNR specific variant read frequency thresholds and allelic bias. The parameters required for classification were determined in a training cohort of 139 CRC tumours where the MSI status had been previously characterised, and a testing cohort of 70 CRC tumours was used for blinded validation of the method.

## Material and methods

### Study samples

The present study consisted of 3 cohort of samples: discovery, training and testing cohorts. The discovery cohort consisted of 132 CRC tumour and tissues samples that were obtained either as formalin fixed paraffin embedded (FFPE) tissues or as DNA extracted from FFPE CRC tissues, from the Northern Genetics Service, Newcastle Hospitals NHS Foundation Trust. The MSI status of all tumours had previously been established using the MSI Analysis System Version 1.2 (Promega, Southampton, UK).

The training cohort consisted of 141 samples which were obtained as extracted DNA from CRC tumours from the Genetics Service of the Complejo Hospitalario de Navarra and the Oncogenetics and Hereditary Cancer Group, IDISNA (Biomedical Research Institute of Navarra, Spain). These samples were used to identify classification parameters. They had previously been tested for MSI using the MSI Analysis System, Version 1.2 (Promega, Southampton, UK). IHC expression analysis was performed using (BD biomedical Tech, New Jersey, USA) antibodies for MLH1 protein at 1:10; MSH6 protein at 1:120; and PMS2 protein at 1:100, and (Oncogene Ltd Middlesex, UK) antibody for MSH2 protein at 1:100 ratio. However, data were available for 124 of the samples only.

The testing cohort consisted of 70 anonymised tumour DNA samples that were obtained from the Department of Molecular Pathology, University of Edinburgh. MMR status had been tested for clinical service use using the MSI Analysis System, Version 1.2 (Promega, Southampton, UK).

The study was conducted following ethical approval from Medical Research and Ethics Committee (CEIC Navarra Government) and Newcastle Hospitals NHS Foundation Trust (REC reference 13/LO/1514).

### *In silico* selection of MNRs

Whole genome sequences consisting of MSI colorectal cancers, matched normals, and MSS cancers were obtained from The Cancer Genome Atlas (TCGA) project (http://cancergenome.nih.gov/; access identifier: phs000178.v8.p7 DAR: 17798, request date 2012-11-13; Study accession phs000544.v1 .p6; parent study: phs000178.v7.p6 ; 35 samples, see S1 Table) [27]. BAM files were converted to Fastq files using bam2fastq (version 1.1.0) [28]. Sequence alignment was performed using BWA (version 0.6.2) [29], indexing and sorting of BAM files was done using samtools (version 0.1.18) [30] and duplicates were removed using PICARD (version 1.75) [31]. GATK (version 2.2.9) [32] was used to produce a combined BAM file for all samples and to realign around indels. The GATK UnifiedGenotyper was used to produce a raw variant call file which was annotated using the TandemRepeatAnnotator for indel identification in mononucleotide repeats. Mononucleotide repeats of lengths 7bp–12bp were selected, and repeats encompassing common sequence variants (dbSNP version 173, hg19) [33] were removed. SNPs listed in dbSNP within 30bp of the repeats were annotated using Perl scripts. Because of the low pass nature of the sequence data, all reads from MSI tumours were combined in one group, while reads from MSS and MSI-L tumours and from normal samples were combined in a second group as controls.

### MNR amplification

Primers were designed using Primer3 [34] or manually if Primer3 returned no suitable oligonucleotides. Primers designed manually had a Tm of 57–60°C. All primers were checked for common SNPs using SNP Check (https://ngrl.manchester.ac.uk/SNPCheckV2/snpcheck.htm), off target binding using BLAST (http://blast.ncbi.nlm.nih.gov/Blast.cgi) or BLAT [35], and appropriate melting temperatures and absence of secondary structures using OligoCalc (http://www.basic.northwestern.edu/biotools/oligocalc.html) or Primer3. Primers were manufactured either by Metabion (Metabion International AG, Steinkirchen, Germany) or by Biobasic (Bio Basic Inc., Markham, Canada). Primers for all MNRs were initially designed to create amplicon of approximately 300–350bp (see S2 Table). For the final MNR panel, a second set of primers were designed to generate 100–150bp amplicons with 5' adapters (see S3 Table). Amplicons were generated using the high fidelity *Pfu*-based Herculase II Fusion DNA polymerase (Agilent, Santa Clara, CA, USA) at 35 PCR cycles following the manufacturer's master mix and PCR cycling condition recommendation recommended by the manufacturer.

### Sequencing

Amplicons were quantified using Qiagen QIAxcel (Qiagen, Manchester UK.), then pooled at roughly equimolar concentrations. Agencourt AMPure XP beads (Beckman-Coulter Life Sciences, Indianapolis, USA) were used for PCR clean up before Library Preparation. For the 300–350bp amplicons, barcoding and library preparation were performed using the Nextera XT DNA Library Prep kit (Illumina, San Diego, CA, United States of America), after pooling of the amplification products for each sample, while for the 100–150 bp amplicons the 16S metagenomic sample preparation protocol was followed (http://support.illumina.com/documents/documentation/chemistry_documentation/16s/16s-metagenomic-library-prep-guide-15044223-b.pdf). Sequencing was performed on the Illumina MiSeq plattform to a target depth of at least 10,000 reads per amplicon per sample.

### Variant and MNR calling

Sequences were aligned using BWA (version 0.6.2) and the hg19 assembly as the reference genome. Samtools was used to sort and index the BAM files, and realignment was done using GATK (3.1.1). Alignment files were converted to SAM format and processed using custom R scripts. Only features observed on both reads of a pair, i.e. concordant in both orientations, were used in subsequent calculations and only amplicons where the MNR was covered by at least 20 read pairs were analysed. Flanking SNPs were considered to be heterozygous if the least common allele, i.e. the allele supported by the smallest number of reads, was present in at least 20% all the read pairs covering the SNP position.

### Construction of MNR specific ROC curves

For each marker, the proportion of reads representing MNR deletion alleles in MSI-H and MSS samples was analysed separately. A threshold approach to instability classification was used: samples with a proportion of variant reads above the threshold being classified as MSI-H and below as MSS. This enabled the relative frequency of true positives (i.e. known MSI-H samples with a value above the threshold), and of false positives (i.e. known MSS samples with a value above the threshold) to be determined. For each MNR, these two values were then plotted against each other for thresholds between 0 and 1. The resulting curve represents the receiver operating characteristic (ROC) curve and the area under the curve (AUC) was used as a quantitative measure of the ability of the MNR to discriminate between MSI-H and MSS samples.

### MNR based classification using deletion frequency and allelic bias

The classifier was designed to include information both on changes in MNR length, and on the distribution of the variant reads across both alleles. Since discrimination between alleles is only possible for samples heterozygous for a flanking SNP, not all samples can be assessed for biased distribution of variant reads across both alleles. However, lack of data should not favour either classification. A naïve Bayes approach for the classification procedure was used [36]. The underlying idea is to compare the probabilities of belonging to one of two classes, i.e. MSI-H or MSS, given the observations at each of the MNR markers used.

If we consider a set of MNRs and, for a particular sample, we represent the observed frequency of reads showing deletion for each of them with *O*, the probability that the sample is microsatellite unstable with *p(MSI|O)*, and the probability that the sample is microsatellite stable with *p(MSI|O)*, then the ratio:

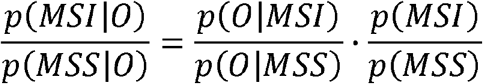

can be used as the discrimination criterion. Here *p(MSI)* and *p(MSS)* designate the *a priori* probability of a sample being MMR deficient or proficient.

An observation consists of the read count data at the different MNRs; i.e.*O = (Oi, …,0*_N_,), where *N* designates the number of MNRs assessed in the assay. Assuming that, for a given mismatch repair status, mutations at the different markers occur independently from each other, then:

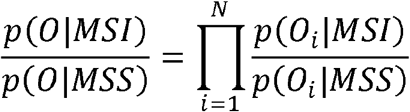

For a microsatellite *i* in each individual, an observation *O_i_* is described by two values, *Di* and *B_t_*, i.e. *O_t_ =* and *p(O_i_) = p(D_i_)p(B_i_\D_i_)*, where = 1 if the number of reads representing a deletion is above a pre-specified threshold and 0 otherwise, and *B_i_* if significant bias was observed and 0 otherwise. Therefore:

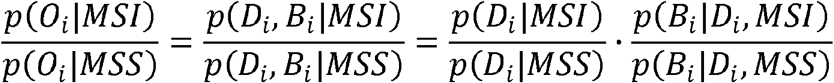

in cases where the bias cannot be computed, for example, when there are no heterozygous flanking polymorphic sites, we set (*O_i_|MSI*) = *p(D_i_|MSI), p(O_i_|MSS) = p(D_i_|MSS)* and the factor 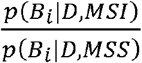 can be omitted.

A threshold for each microsatellite was chosen, such that 95% of all MSS samples have frequencies below the threshold. To estimate and *p(D_i_|MSI)*, the exact numbers of MSS and MSI-H samples with frequencies above the threshold were used. To estimate *p(B_i_|D_i_,MSS)* and *p(B_i_|D_if_MSS*), samples heterozygous at a flanking SNP marker, and for which the frequency of reads with deletions exceeded the MNR specific thresholds, were used. Bias was considered to be present when the association between the presence of a deletion and the genotype at the flanking SNP was significant at p-value of 0.05 using Fishers' exact test. If there were multiple heterozygous SNPs neighbouring a repeat then the SNP with the lowest p-value was used. When the deletion frequency was below the threshold, *p(B_i_|D_i_,MSI)* and *p(B_i_|Di,MSS)* were set to 1. This is equivalent to assuming that in such cases there is insufficient evidence for an MNR mutation and therefore bias is not meaningful.

The results are presented as a score 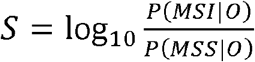

Here we used a set of samples to determine, for each MNR, the following parameters used in the classification: (1) A threshold for the frequency of reads showing a deletion (for the choice of thresholds see previous paragraph and the results section for an illustration); (2) The proportion of MSI-H samples with a deletion frequency above this threshold; (3) The proportion of MSS samples with a deletion frequency above the threshold; (4) The proportion of MSI-H samples showing a deletion and significant allelic imbalance and; (5) The proportion of MSS samples showing a deletion and significant allelic imbalance. The frequencies of MSS and MSI-H tumours were assumed to be 0.85 and 0.15 respectively [37], i.e. *p(MSS) = 0.85* and *p(MSI) = 0.15*.

These parameters were then used to calculate the score for each tumour in a second, independent set of samples. Samples with a score below 0 were classified as MSS and those above as MSI-H.

## Results

### Identification of MNR panel

A total of 218,181 variable 7–12bp MNRs were identified from the TGCA CRC genome sequence data. From these, we excluded MNRs with a read depth less than 20 in either the MSI-H or the MSS group, and MNRs that did not have a SNP (dbSNP137) with a minor allele frequency larger than 20% within 30bp of the repeat. MNRs with multiple lower frequency SNPs in the flanking regions were not excluded if the probability of observing a minor allele in at least one SNP was above 20%, assuming linkage equilibrium.

For MNRs with a length of 7–9bp, only those, which had no length variation in the control group but where at least 10% of reads in the MSI-H group showed length variants, were selected. For MNRs with a length of 10–12bp, only MNRs where the frequency of reads showing length variation was at most 5% among controls and at least 15% among MSI-H samples, were selected. In total, 529 poly A-MNRs fulfilled these criteria. For poly C-MNRs no microsatellite fulfilled these criteria. To ensure that some polyC MNRs were included in subsequent analyses, the minimum depth and flanking SNPs requirements were dropped, leading to the selection of 33 polyC MNRs. From these 562 markers, MNRs within repetitive elements and regions of low complexity (likely to be refractory to amplicon design) were also excluded, producing a final list of 120 MNRs (S2 Table).

To eliminate potentially uninformative repeats, amplicons were designed for all 120 MNRs. These were initially tested in FFPE samples from the discovery cohort: 6 tumours from patients with Lynch syndrome, 5 normal mucosa samples and 6 samples from sporadic MSS tumours. Amplicons were pooled, indexed, and sequenced to a target depth of 10,000 reads per amplicon per sample. Only results for amplicons represented by at least 100 paired end reads were analysed and representative results are shown in Fig 1.

**Fig 1:**
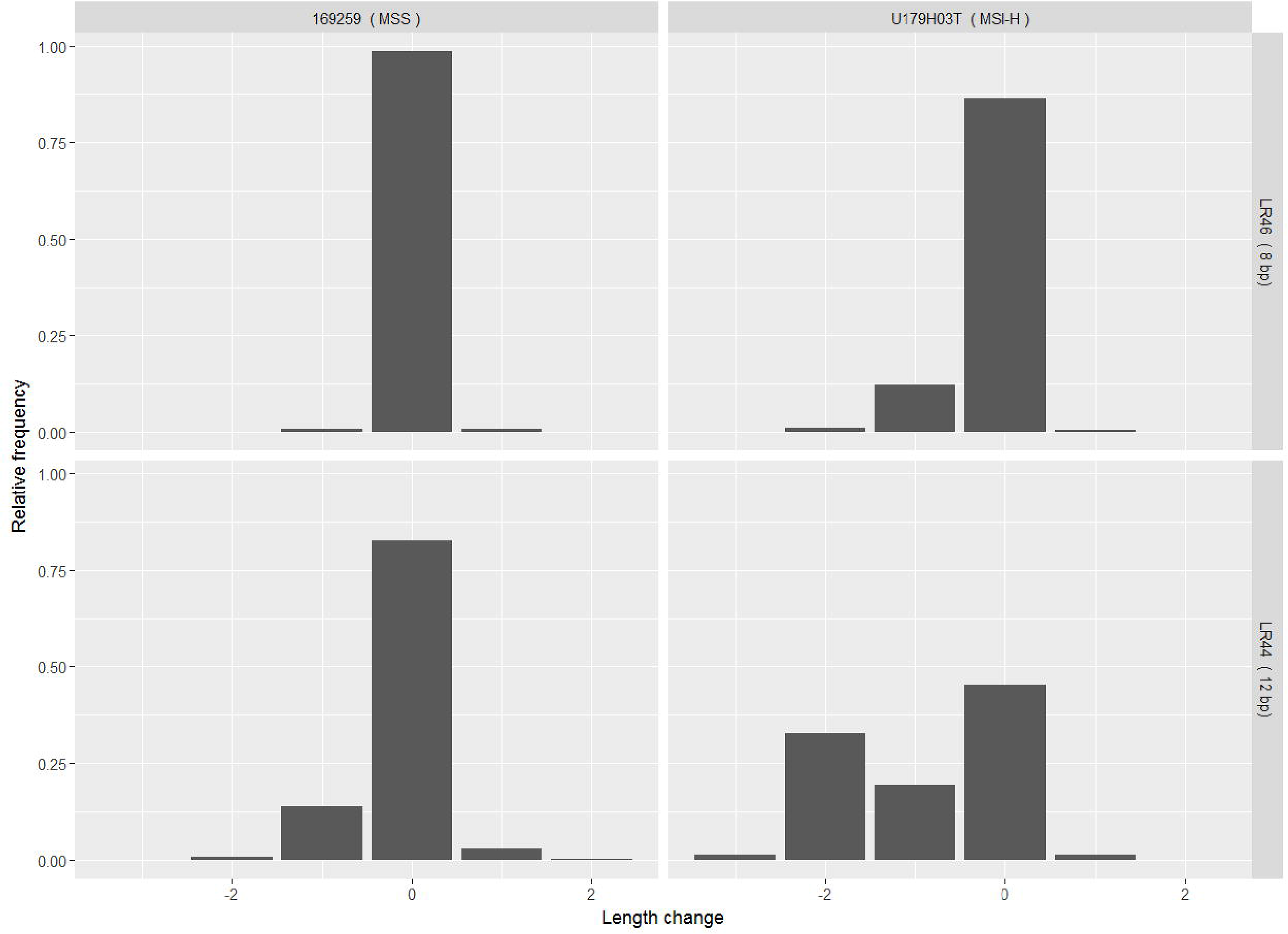
Example distributions of read frequencies. Relative frequencies of reads classified according to length are shown for MNRs LR46 (an 8bp long poly-A tract) and LR44 (12 bp poly-A) in an MSS sample (169259) and MSI-H sample (U179H03T). The abscissa represents the deviation from the reference sequence length in bp.

Fig 1 shows the relative frequencies of reads for two MNRs in an MMR proficient (MSS) and an MMR deficient (MSI-H) sample. A small fraction of insertion reads (+1 value in the abscissa) are observed in both MSI-H and MSS samples, but the frequency of deletions (−1, −2 and −3 values) differs between the two. However, for the longer repeat shown, reads representing deletions of more than one base pair are also observed in the MSS sample, while a second peak can be observed corresponding to a 2 bp deletion in the MSI-H sample. In all analyses, the sum of the frequencies of reads representing all deletions were used.

To illustrate levels of allelic variation observed, results from a single MNR marker (LR46) are shown in Fig 2. The read distribution for each allele is plotted separately for an MSI-H and an MSS sample that are heterozygous for the flanking SNP. While the distributions for both the G and A alleles in the MSS sample are similar, reads representing a one base pair deletion are predominantly found in the G allele of the MSI-H sample.

**Fig 2:**
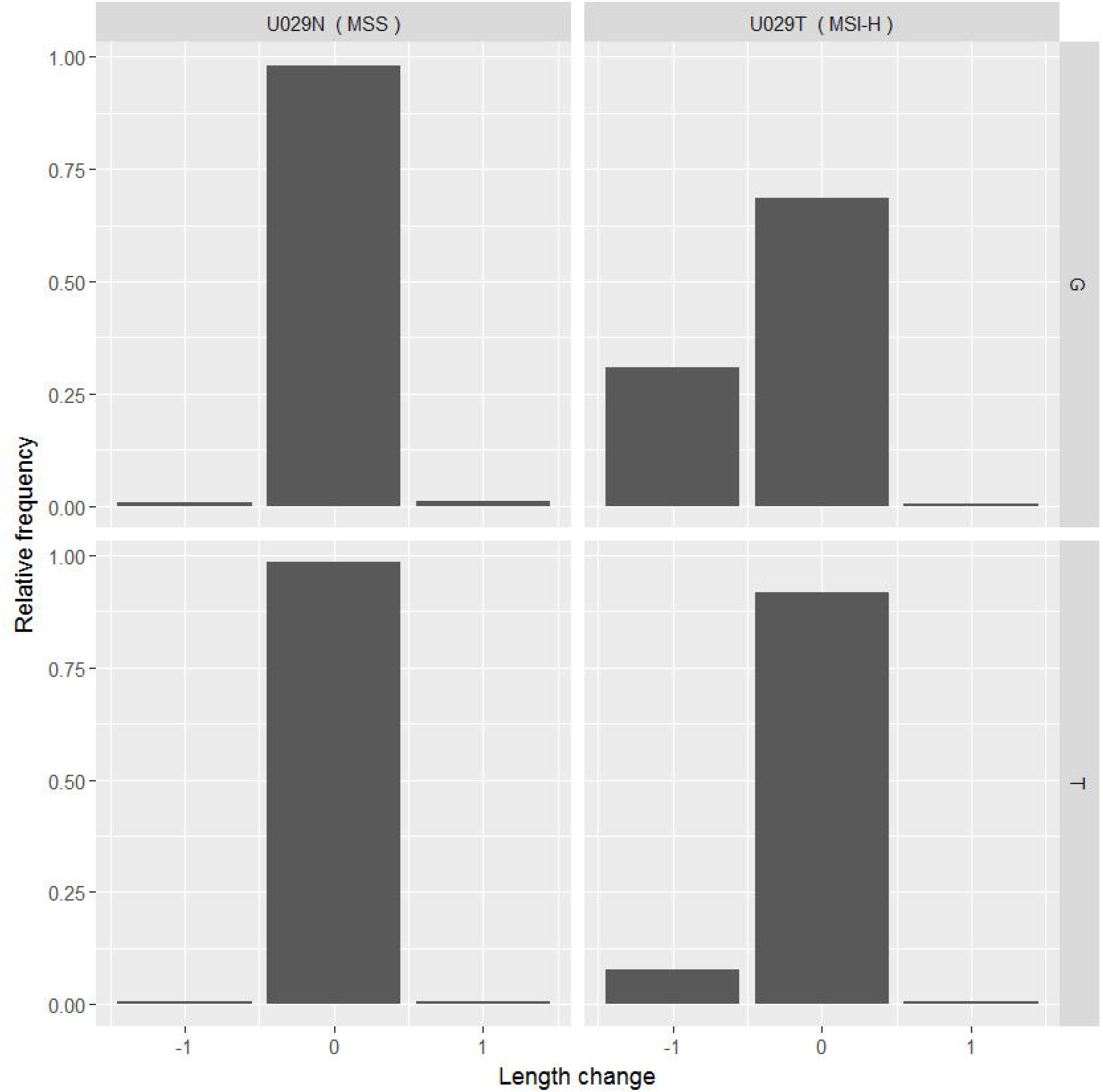
Example representing allelic bias. Allele specific read frequencies and sizes are shown for LR46 in two samples from a patient who is heterozygous for a flanking SNP (rs6040079). U029N=normal somatic tissue, U029T=microsatellite unstable tumour.

From this initial assessment, MNRs were retained for further analysis only if they exhibited a deletion frequency >5% in 1 or more MSI-H sample, and these frequencies were also >1.5 fold higher than frequencies observed in all normal mucosa samples. 41 MNRs satisfied these criteria. Two previously described MNRs adjacent to SNPs (one in DEPDC2 [38] and one in the intergenic repeat AP0035322 [39]) were also added to the analysis at this stage. These 43 MNRs were each typed in a minimum of 28 MSI-H and 30 MSS tumours in the discovery cohort, and ROC curves were generated to assess the ability of each to discriminate between MSI-H and MSS samples. This was performed by estimating the area under the curve (AUC) using the frequency of reads representing MNR deletion as the classification criterion, and classifying samples with a frequency above each threshold as MSI-H and below each threshold as MSS (S2 Table).

Representative examples of this analysis are shown in Fig 3 which shows the ROC curves for the two poly-A MNRs-LR46 (8bp) and LR44 (12bp) used in Fig 1. The AUC for LR46 was 0.83 (95% confidence interval 0.71–0.84) and 0.99 (0.98 0. 99) for LR44. Using the AUC and amplicon length as a criterion, 15 poly-A MNR repeats were selected and together with the two poly-C MNR with the largest AUC formed our final panel (S3 Table). As described in the Methods section, the primers for this panel were redesigned to produce shorter amplicons (S3 Table).

**Fig 3:**
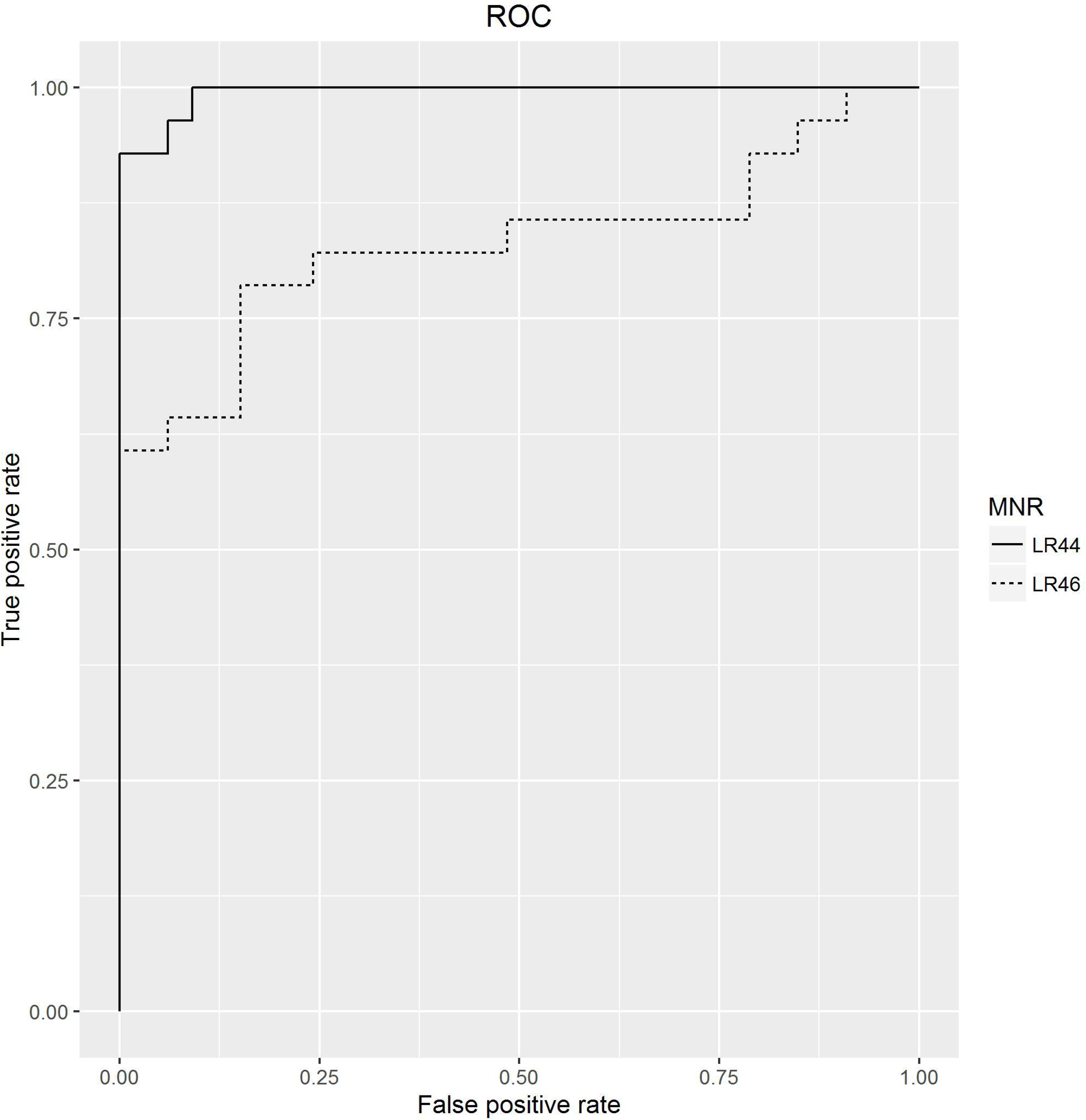
Assessing the ability of single MNRs to discriminate between MMR proficient and deficient samples.

### Tumour classification using the selected panel of short MNRs

To establish the parameters required by the classification procedure, the seventeen MNRs included in the final panel were typed in the training cohort consisting of 139 samples, of which 67 had been classified as MSI-H by fragment analysis. The deletion frequencies and allelic biases observed in these samples were used to establish thresholds for each marker and to estimate the probabilities described in the methods section for MSI-H and MSS samples. To illustrate this step, results for LR44, a 12 bp poly-A MNR, are presented in Fig 4. Fig 4A depicts the distribution of the relative frequencies of reads showing deletions in LR44. As expected, the deletion frequency is higher in MSI-H tumours. The horizontal line represents a threshold of 0.24 (see Methods section for the choice of threshold). The deletion frequency was higher than the threshold in 58 of the 66 MSI-H samples for which data were available for this marker, but only in 4 of the 72 MSS samples.

**Fig 4:**
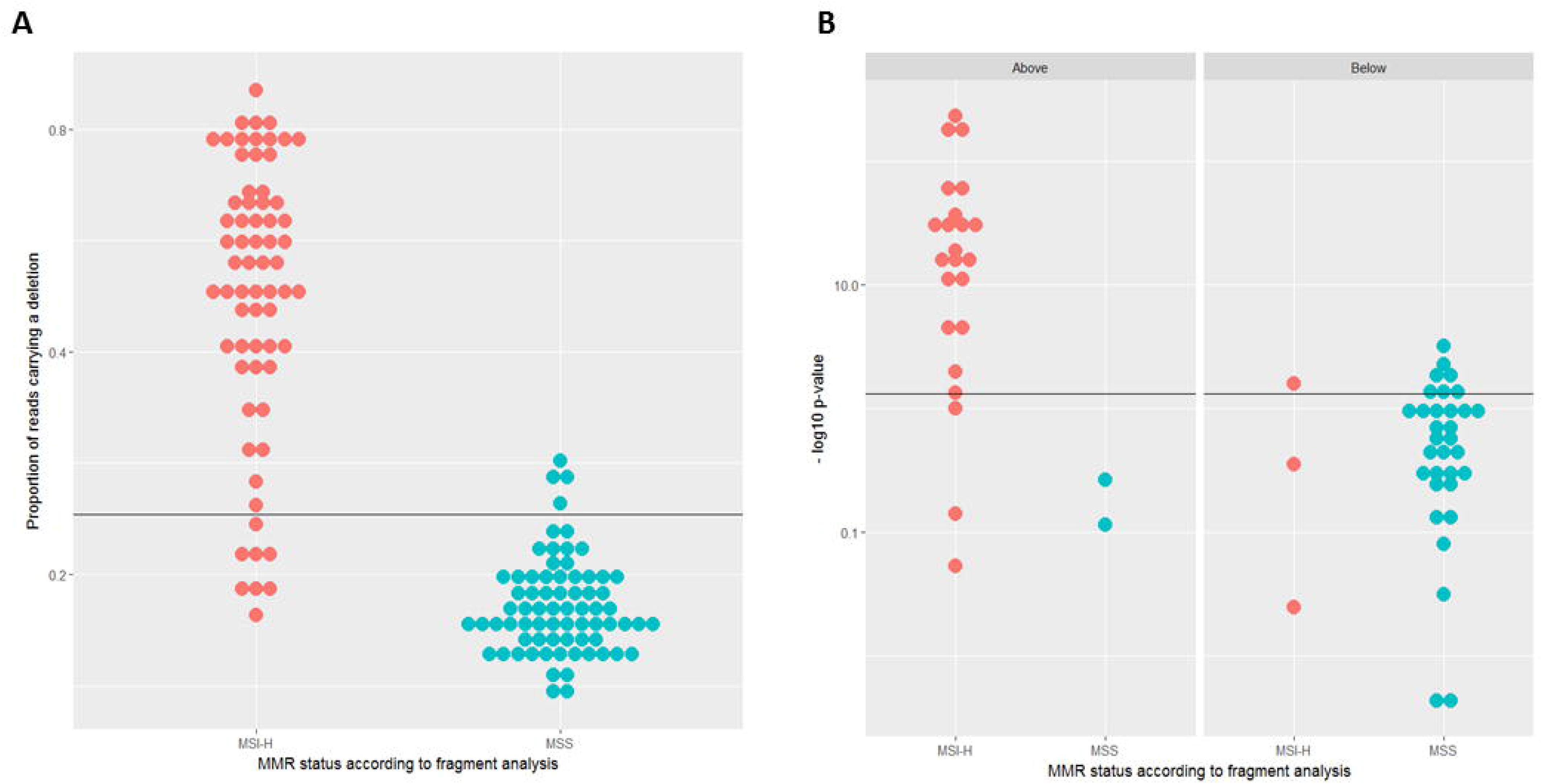
Establishing analysis parameters for MSI test. (A) Relative frequency of reads carrying a deletion in MSI-H and MSS samples for the MNR LR44. (B) Analysis of allelic bias at the MNR LR44 for MSI-H and MSS samples stratified according to the proportion of reads showing deletions. MSI-H= Microsatellite instability-high. MSS= Microsatellite stable.

Of the 139 samples depicted in Fig 4A, 60 samples (26 MSI-H and 34 MSS) were heterozygous for a SNP flanking the repeat, and the distribution of allelic bias for these samples is presented in Fig 4B. Fisher's exact test was used to assess whether deletion reads were evenly distributed between both alleles. The Figure represents the resulting p-values -log_10_(*p*) in a scale. The left hand panel shows the heterozygous samples that are above the threshold in Fig 4A, the right hand panel those that are below. Overall, 21 MSI-H and 4 MSS samples had values above the threshold, i.e. had a bias significant at the 5% level. This corresponds to our expectation that allelic bias will be more common among MSI-H samples.

It is noteworthy that only 2 of the 4 MSS samples above the frequency threshold in Fig 4A were heterozygous, and neither showed significant bias. In contrast, 27 out of the 32 MSI-H samples which were heterozygous showed a bias above the threshold (Fig 4B). This difference is significant (two sided Fishers' exact p=0.03), while the corresponding test for samples that do not reach the frequency threshold does not suggest any difference between MSS and MSI-H samples (two sided Fishers' exact p=0.39). This is consistent with our assumption that allelic bias can help to discriminate between MSI-H and MSS samples. For allelic bias and deletion frequencies, thresholds and relative numbers of samples above the respective threshold were determined for each of the 17 MNRs.

The parameters determined in the training cohort were then used to test the procedure in an independent testing cohort consisting of 70 CRC samples, 36 of which had previously been classified as MSI-H and 34 as MSS. Fig 5 presents the contribution made to tumour classification by MNR length variation (Fig 5A) and MNR allelic bias (Fig 5B). This illustrates that while both contribute to the separation of the groups; changes in MNR length provide the main contribution. The final combined classification (Fig 5C) is concordant with fragment analysis, achieving 100% sensitivity and specificity (95% confidence intervals 87–100% and 90–100%, respectively) when fragment analysis is used as the reference technique.

**Fig 5:**
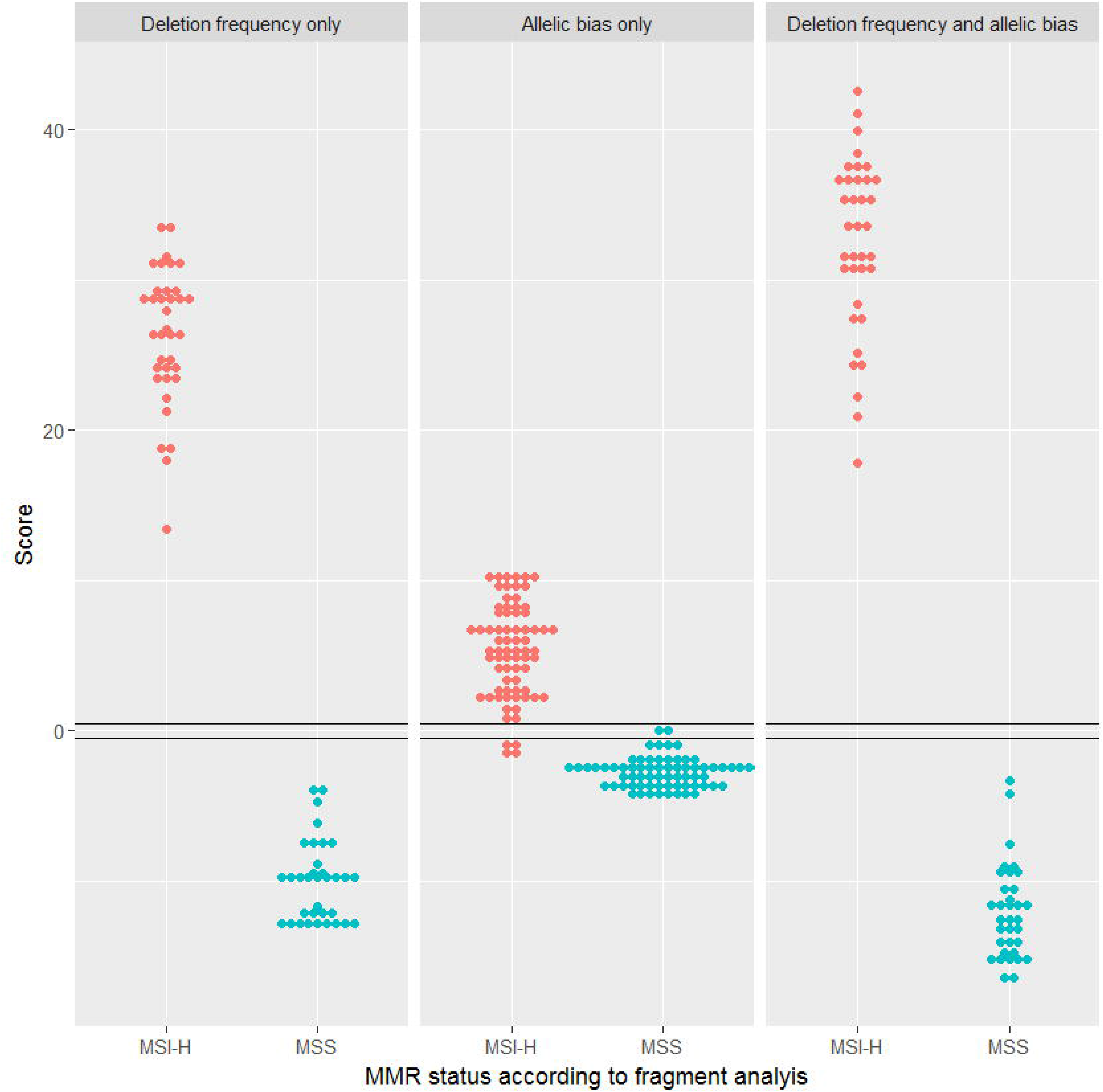
Classification of MMR status in an independent dataset of 70 CRC samples. Classification using only deletion frequency data (A), only allelic bias data (B), and both parameters combined (C). MSI-H= Microsatellite instability-high. MSS= Microsatellite stable.

Finally, we used the data from the testing cohort to estimate the parameters and classify the samples in the training cohort. The results are represented in Fig 6. Four samples gave discordant results relative to fragment analysis (samples 63, 72, 91 and 135). Immunohistochemistry for sample 63 was checked and found to be consistent with reported MSS status. However, DNA from sample 72 was reanalysed by fragment analysis and MSI-H phenotype was detected, while IHC analysis of samples 91 and 135 revealed no alteration in expression for *MSH2, MLH1, MSH6* and *PMS2* genes. This raises the possibility that IHC and fragment analysis are inconsistent for these 3 samples. Overall, there was a 92% concordance between fragment analysis and IHC, as assessed by staining for MSH2, MLH1, MSH6 and PMS2 proteins. For this analysis, the concordance between our results and fragment analysis is 97% and the estimates for sensitivity and specificity are both 97% (95% CI: 89–99% and 90–99%, respectively) when results from fragment analysis are used as reference. Interestingly, reclassification using the training cohort for both parameter estimation and for testing the classification resulted in misclassification of the same four samples. Combining both sets of results led to a sensitivity of 98% (95% CI: 92–99%) and specificity of 98% (95% CI: 93–99%).

**Figure 6:**
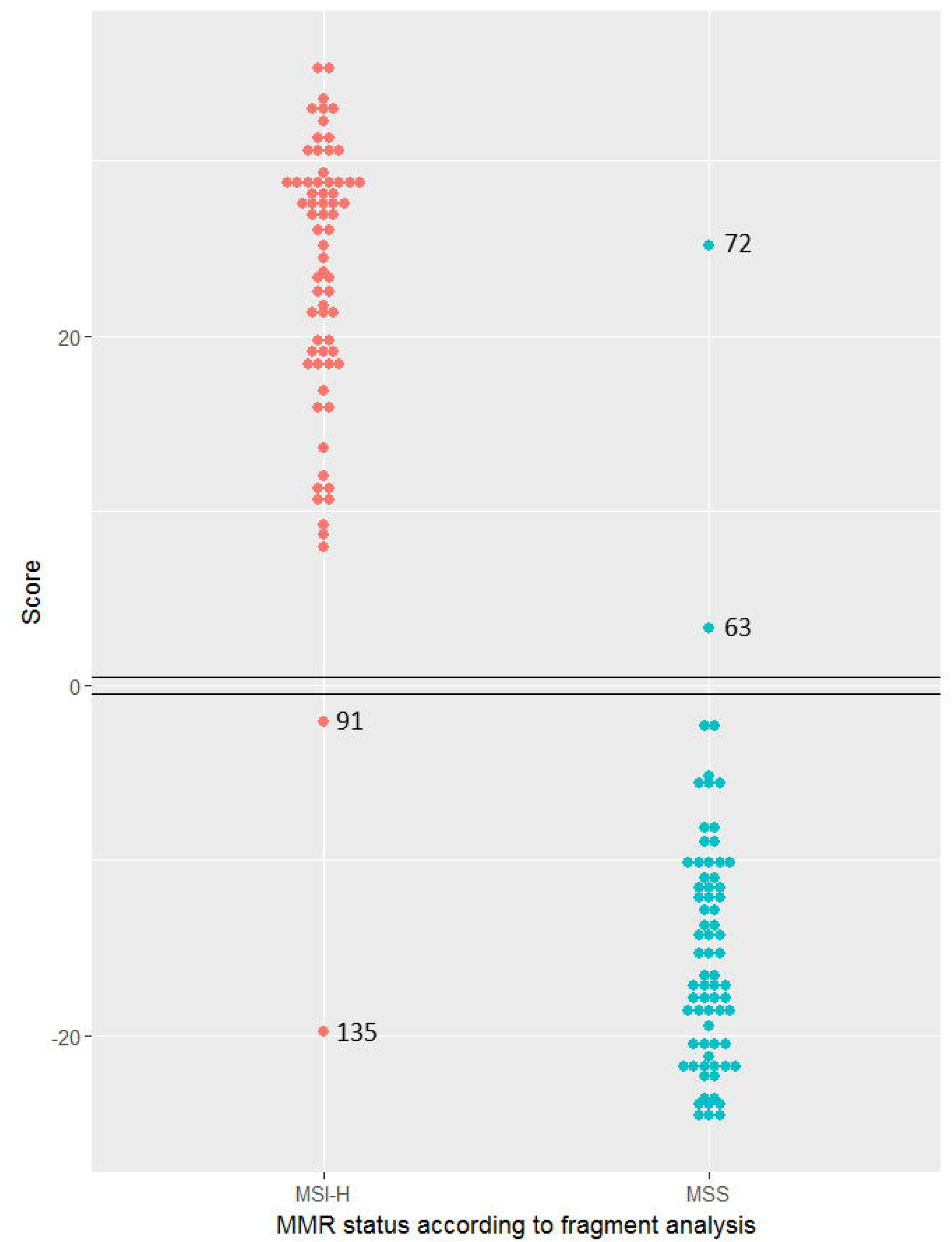
MMR status classification of the training set. MSI-H= Microsatellite instability-high. MSS= Microsatellite stable.

## Discussion

The method presented here allows sequence-based discrimination between MSI-H and MSS tumours using a limited number of loci, without the requirement for paired germline DNA as a reference. A multi-step process was used to select a panel of MNRs involving analysis of genomic sequence data to identify the most promising markers, and two rounds of amplicon assessment. Although, this does not ensure that the optimal set of MNRs was identified, the performance of the panel is comparable to that of fragment analysis.

We chose relatively short MNRs for our test to diminish the probability of PCR artefacts and to reduce the likelihood of encountering germline variation affecting MNR length, a potential confounding factor in cases where no normal material is available. However, somatic instability is also lower meaning that genuine mutations will tend to affect only one allele. Therefore, even allowing for PCR errors, mutant reads should concentrate on one allele. We showed that this can be assessed using flanking heterozygous SNPs and can be used to improve classification. It is worth noting that even in situations where mutations have occurred in both alleles, each allele is likely to be affected in a different proportion of cells in a sample since, during clonal evolution, there will be a time interval between the occurrence of the two mutations, and this time interval is expected to be larger for shorter microsatellites.

To our knowledge, this is the first method for assessing MSI that uses allelic information. Although we only use allelic data to assess bias in the distribution of mutant reads, it can also help to distinguish between somatic and germline variation, in particular in situations where no normal material is available, but the tumour is expected to contain normal tissue contamination. MNRs showing germline variants can be excluded from the analysis although it would also be possible to treat each allele separately. Allelic analysis, however, is only possible for MNRs heterozygous for flanking SNPs in a particular sample. In principle, it would be feasible to restrict the score calculation to such MNRs. However, such a procedure would disregard information from many of the amplicons used, and require larger marker panels, increasing assay costs.

Here we used thresholds on the frequency of reads representing mutated MNRs because we wanted to dichotomise the data. Other approaches would be possible, however, using a threshold that is above the frequency observed in the majority of the MSS samples is consistent with the approaches followed by other authors who aim to set their thresholds so that variation reflecting PCR artefacts is excluded [24]. The formalism presented here could be used without defining thresholds, but this would require specifying the whole deletion frequency distribution. Similarly, we used a threshold, the p-value of 0.05 in Fisher's exact test, to dichotomise allelic bias. Using the statistical significance of the bias seems natural although the precise choice of the threshold is arbitrary.

Since our test aims to detect MSI-H tumours, it seems reasonable to use fragment analysis as the reference technique. However, MSI detection is usually a means for assessing MMR proficiency. It is noteworthy that in 3 out of 4 cases where there were discrepancies between our results and the results from fragment analysis, there were also discrepancies between fragment analysis and IHC results.

In summary, we propose an approach to the detection of MSI-H tumours whose main advantage is its simplicity, making it suitable for high throughput analysis without the need for control normal DNA. Establishing whether tumours have resulted from a breakdown in mismatch repair is important in clinical management of the individual and can help prevent future cancers in those families where there is a germline molecular defect. Expansion of testing to all colorectal cancers has been shown to be cost effective in the UK [40] and is soon to become standard of care on the basis of National Institute of Healthcare and Clinical Excellence (NICE) guidance [DG27] in the UK's National Health Service [41]. Similar decisions are being taken in other developed nations. A scalable, reliable MSI test will have clinical utility while modest costs and the ability to link this analysis to routine pathology assessment with help to ensure rapid adoption and facilitate further molecular approaches to tumour profiling and precision medical care.

## Supporting information

S1 Table: Sample identifiers for whole genome sequences obtained from The Cancer genome Atlas (TCGA) database used in this study.

S2 Table: List containing 300–500bp amplicon or repeat name of 120 markers, amplicon position (genome build hg19), PCR primer sequences, SNPs in close proximity to mononucleotide repeats and AUC of 41 selected markers.

S3 Table: List containing amplicon or repeat name (100–150bp) of 17 marker panel, amplicon position (genome build hg19), PCR primer sequences and SNPs in close proximity to mononucleotide repeats.

## References

1. Ionov Y, Peinado MA, Malkhosyan S, Shibata D, Perucho M. Ubiquitous somatic mutations in simple repeated sequences reveal a new mechanism for colonic carcinogenesis. Nature. 1993;363(6429):558–61.

2. Tran HT, Keen JD, Kricker M, Resnick MA, Gordenin DA. Hypermutability of homonucleotide runs in mismatch repair and DNA polymerase proofreading yeast mutants. Molecular and Cellular Biology. 1997;17(5):2859–65.

3. Palomaki GE, McClain MR, Melillo S, Hampel HL, Thibodeau SN. EGAPP supplementary evidence review: DNA testing strategies aimed at reducing morbidity and mortality from Lynch syndrome. Genet Med. 2009;11 (1):42–65. doi: 10.1097/GIM.0b013e31818fa2db. PubMed PMID: 19125127; PubMed Central PMCID: PMC2743613.

4. Guindalini RS, Win AK, Gulden C, Lindor NM, Newcomb PA, Haile RW, et al. Mutation spectrum and risk of colorectal cancer in African American families with Lynch syndrome. Gastroenterology. 2015;149(6):1446–53. doi: 10.1053/j.gastro.2015.07.052. PubMed PMID: 26248088; PubMed Central PMCID: PMC4648287.

5. Engel C, Loeffler M, Steinke V, Rahner N, Holinski-Feder E, Dietmaier W, et al. Risks of less common cancers in proven mutation carriers with lynch syndrome. J Clin Oncol. 2012;30(35):4409–15. doi: 10.1200/JC0.2012.43.2278. PubMed PMID: 23091106.

6. Popat S, Hubner R, Houlston RS. Systematic review of microsatellite instability and colorectal cancer prognosis. J Clin Oncol. 2005;23(3):609–18. Epub 2005/01/22. doi: 10.1200/jco.2005.01.086. PubMed PMID: 15659508.

7. Le DT, Uram JN, Wang H, Bartlett BR, Kemberling H, Eyring AD, et al. PD-1 blockade in tumors with mismatch-repair deficiency. New England Journal of Medicine. 2015;372(26):2509–20.

8. Hampel H, Frankel WL, Martin E, Arnold M, Khanduja K, Kuebler P, et al. Feasibility of screening for Lynch syndrome among patients with colorectal cancer. J Clin Oncol. 2008;26(35):5783–8.

9. Burn J, Gerdes AM, Macrae F, Mecklin JP, Moeslein G, Olschwang S, et al. Long-term effect of aspirin on cancer risk in carriers of hereditary colorectal cancer: an analysis from the CAPP2 randomised controlled trial. Lancet. 2011 ;378(9809):2081–7. Epub 2011/11/01. doi: 10.1016/s0140-6736(11)61049-0. PubMed PMID: 22036019; PubMed Central PMCID: PMCPMC3243929.

10. Vasen HF, Blanco I, Aktan-Collan K, Gopie JP, Alonso A, Aretz S, et al. Revised guidelines for the clinical management of Lynch syndrome (HNPCC): recommendations by a group of European experts. Gut. 2013;62(6):812–23.

11. Canard G, Lefevre JH, Colas C, Coulet F, Svrcek M, Lascols O, et al. Screening for Lynch syndrome in colorectal cancer: are we doing enough? Ann Surg Oncol. 2012;19(3):809–16.

12. Mills AM, Liou S, Ford JM, Berek JS, Pai RK, Longacre TA. Lynch syndrome screening should be considered for all patients with newly diagnosed endometrial cancer. Am J Surg Pathol. 2014;38(11):1501–9. Epub 2014/09/18. doi: 10.1097/pas.0000000000000321. PubMed PMID: 25229768.

13. Julie C, Tresallet C, Brouquet A, Vallot C, Zimmermann U, Mitry E, et al. Identification in daily practice of patients with Lynch syndrome (hereditary nonpolyposis colorectal cancer): revised Bethesda guidelines-based approach versus molecular screening. The American journal of gastroenterology. 2008;103(11):2825–35; quiz 36. Epub 2008/09/02. doi: 10.1111/j.1572-0241.2008.02084.x. PubMed PMID: 18759827.

14. de la Chapelle A, Hampel H. Clinical relevance of microsatellite instability in colorectal cancer. Journal of Clinical Oncology. 2010;28(20):3380–7.

15. Laiho P, Launonen V, Lahermo P, Esteller M, Guo M, Herman JG, et al. Low-level microsatellite instability in most colorectal carcinomas. Cancer Res. 2002;62(4):1166–70. PubMed PMID: 11861399.

16. Boyle TA, Bridge JA, Sabatini LM, Nowak JA, Vasalos P, Jennings LJ, et al. Summary of microsatellite instability test results from laboratories participating in proficiency surveys: proficiency survey results from 2005 to 2012. Arch Pathol Lab Med. 2014;138(3):363–70.

17. Shinde D, Lai Y, Sun F, Arnheim N. Taq DNA polymerase slippage mutation rates measured by PCR and quasi likelihood analysis:(CA/GT) n and (A/T) n microsatellites. Nucleic acids research. 2003;31(3):974–80.

18. Umar A, Boland CR, Terdiman JP, Syngal S, de la Chapelle A, Ruschoff J, et al. Revised Bethesda Guidelines for hereditary nonpolyposis colorectal cancer (Lynch syndrome) and microsatellite instability. Journal of the National Cancer Institute. 2004;96(4):261–8.

19. Shia J. Immunohistochemistry versus microsatellite instability testing for screening colorectal cancer patients at risk for hereditary nonpolyposis colorectal cancer syndrome. Part I. The utility of immunohistochemistry. The Journal of molecular diagnostics: JMD. 2008;10(4):293–300. doi: 10.2353/jmoldx.2008.080031. PubMed PMID: 18556767; PubMed Central PMCID: PMC2438196.

20. Zhang L. Immunohistochemistry versus microsatellite instability testing for screening colorectal cancer patients at risk for hereditary nonpolyposis colorectal cancer syndrome. Part II. The utility of microsatellite instability testing. The Journal of molecular diagnostics: JMD. 2008;10(4):301–7. doi: 10.2353/jmoldx.2008.080062. PubMed PMID: 18556776; PubMed Central PMCID: PMC2438197.

21. Niu B, Ye K, Zhang Q, Lu C, Xie M, McLellan MD, et al. MSIsensor: microsatellite instability detection using paired tumor-normal sequence data. Bioinformatics. 2014;30(7):1015–6.

22. Lu Y, Soong TD, Elemento O. A novel approach for characterizing microsatellite instability in cancer cells. PLoS One. 2013;8(5):e63056.

23. Zhu L, Huang Y, Fang X, Liu C, Deng W, Zhong C, et al. A Novel and Reliable Method to Detect Microsatellite Instability in Colorectal Cancer by Next-Generation Sequencing. The Journal of molecular diagnostics : JMD. 2018;20(2):225–31. Epub 2017/12/27. doi: 10.1016/j.jmoldx.2017.11.007. PubMed PMID: 29277635.

24. Salipante SJ, Scroggins SM, Hampel HL, Turner EH, Pritchard CC. Microsatellite instability detection by next generation sequencing. Clin Chem. 2014;60(9):1192–9. Epub 2014/07/06. doi: 10.1373/clinchem.2014.223677. PubMed PMID: 24987110.

25. Susanti S, Fadhil W, Ebili HO, Asiri A, Nestarenkaite A, Hadjimichael E, et al. N_LyST: a simple and rapid screening test for Lynch syndrome. Journal of clinical pathology. 2018. Epub 2018/02/24. doi: 10.1136/jclinpath-2018-205013. PubMed PMID: 29472252.

26. Ananda G, Walsh E, Jacob KD, Krasilnikova M, Eckert KA, Chiaromonte F, et al. Distinct mutational behaviors differentiate short tandem repeats from microsatellites in the human genome. Genome Biol Evol. 2013;5(3):606–20. doi: 10.1093/gbe/evs116. PubMed PMID: 23241442; PubMed Central PMCID: PMC3622297.

27. Cancer Genome Atlas N. Comprehensive molecular characterization of human colon and rectal cancer. Nature. 2012;487(7407):330–7. doi: 10.103B/nature11252. PubMed PMID: 22810696; PubMed Central PMCID: PMC3401966.

28. bam2fastq software [http://gsl.hudsonalpha.org/information/software/bam2fastq].

29. Li H, Durbin R. Fast and accurate short read alignment with Burrows-Wheeler transform. Bioinformatics. 2009;25(14):1754–60. Epub 2009/05/20. doi: 10.1093/bioinformatics/btp324. PubMed PMID: 19451168; PubMed Central PMCID: PMCPmc2705234.

30. Li H, Handsaker B, Wysoker A, Fennell T, Ruan J, Homer N, et al. The Sequence Alignment/Map format and SAMtools. Bioinformatics. 2009;25(16):2078–9. Epub 2009/06/10. doi: 10.1093/bioinformatics/btp352. PubMed PMID: 19505943; PubMed Central PMCID: PMCPmc2723002.

31. PICARD [http://picard.sourceforge.net].

32. DePristo MA, Banks E, Poplin R, Garimella KV, Maguire JR, Hartl C, et al. A framework for variation discovery and genotyping using next-generation DNA sequencing data. Nature genetics. 2011; 43(5):491–8. Epub 2011/04/12. doi: 10.103B/ng.806. PubMed PMID: 2147BBB9; PubMed Central PMCID: PMCPmc30B3463.

33. Sherry ST, Ward MH, Kholodov M, Baker J, Phan L, Smigielski EM, et al. dbSNP: the NCBI database of genetic variation. Nucleic acids research. 2001 ;29(1):308–11. Epub 2000/01/11. PubMed PMID: 11125122; PubMed Central PMCID: PMCPmc297B3.

34. Rozen S, Skaletsky H. Primer3 on the WWW for general users and for biologist programmers. Methods in molecular biology (Clifton, NJ). 2000;132:365–86. Epub 1999/11/05. PubMed PMID: 10547847.

35. Kent WJ. BLAT-the BLAST-like alignment tool. Genome Res. 2002;12(4):656–64. Epub 2002/04/05. doi: 10.1101/gr.229202. Article published online before March 2002. PubMed PMID: 11932250; PubMed Central PMCID: PMCPmc187518.

36. Gelman A. Bayesian data analysis. Third edition. ed. Boca Raton: CRC Press; 2014. xiv, 661 pages p.

37. Boland CR, Goel A. Microsatellite instability in colorectal cancer. Gastroenterology. 2010;138(6):2073–87e3. doi: 10.1053/j.gastro.2009.12.064. PubMed PMID: 20420947; PubMed Central PMCID: PMC3037515.

38. Alhopuro P, Phichith D, Tuupanen S, Sammalkorpi H, Nybondas M, Saharinen J, et al. Unregulated smooth-muscle myosin in human intestinal neoplasia. Proceedings of the National Academy of Sciences of the United States of America. 2008;105(14):5513–8. Epub 2008/04/09. doi: 10.1073/pnas.0801213105. PubMed PMID: 18391202; PubMed Central PMCID: PMCPmc2291082.

39. Sammalkorpi H, Alhopuro P, Lehtonen R, Tuimala J, Mecklin JP, Jarvinen HJ, et al. Background mutation frequency in microsatellite-unstable colorectal cancer. Cancer Res. 2007;67(12):5691–8. Epub 2007/06/19. doi: 10.1158/0008-5472.can-06-4314. PubMed PMID: 17575135.

40. Snowsill T, Huxley N, Hoyle M, Jones-Hughes T, Coelho H, Cooper C, et al. A systematic review and economic evaluation of diagnostic strategies for Lynch syndrome. Health Technol Assess. 2014;18(58):1–406.

41. NICE. Molecular testing strategies for Lynch syndrome in people with colorectal cancer 2017 [cited 2017 18/04/2017]. Available from: https://www.nice.org.uk/guidance/dg27/chapter/1-Recommendations.

